# The chemical Langevin equation for biochemical systems in dynamic environments

**DOI:** 10.1101/2021.12.19.473404

**Authors:** Lucy Ham, Megan A. Coomer, Michael P. H. Stumpf

## Abstract

Modelling and simulation of complex biochemical reaction networks form cornerstones of modern biophysics. Many of the approaches developed so far capture temporal fluctuations due to the inherent stochasticity of the biophysical processes, referred to as intrinsic noise. Stochastic fluctuations, however, predominantly stem from the interplay of the network with many other — and mostly unknown — fluctuating processes, as well as with various random signals arising from the extracellular world; these sources contribute extrinsic noise. Here we provide a computational simulation method to probe the stochastic dynamics of biochemical systems subject to both intrinsic and extrinsic noise. We develop an extrinsic chemical Langevin equation—a physically motivated extension of the chemical Langevin equation— to model intrinsically noisy reaction networks embedded in a stochastically fluctuating environment. The extrinsic CLE is a continuous approximation to the Chemical Master Equation (CME) with time-varying propensities. In our approach, noise is incorporated at the level of the CME, and can account for the full dynamics of the exogenous noise process, irrespective of timescales and their mismatches. We show that our method accurately captures the first two moments of the stationary probability density when compared with exact stochastic simulation methods, while reducing the computational runtime by several orders of magnitude. Our approach provides a method that is practical, computationally efficient and physically accurate to study systems that are simultaneously subject to a variety of noise sources.

## I. INTRODUCTION

Random fluctuations of molecule numbers play a significant and often functional role at the single cell level. This biochemical noise has been shown to be responsible for a wide variety of observed phenomena. Cell-fate decision making^1–3^, non-genetic heterogeneity^4,5^ and state-switching in dynamic environments or phenotypic plasticity^6–10^ are all examples that are underpinned by stochasticity of the sub-cellular molecular reactions. We therefore require modelling approaches that are able to account for these intricate and sometimes counter-intuitive stochastic effects. The cornerstone for studying stochastic biochemical kinetics remains the stochastic simulation algorithm (SSA)^11^.

The introduction of noise makes *in-silico* simulations computationally expensive. Since Monte Carlo schemes such as the SSA execute each and every reaction event in a system, and require multiple sample trajectories to back calculate the time evolution of the probability density function, these issues quickly render the simulation of biologically relevant models intractable. As a result, significant effort has been expended on the development of approximation methods for solving stochastic systems, and a large variety of different methods have emerged^12–16^. A prominent approximation method is the chemical Langevin equation (CLE)^17^, which can be thought of as a stepping stone between the SSA and the conventional deterministic reaction rate equation^18^. In this intermediate regime, stochasticity is still important, but there exist a sufficient number of molecules to describe the kinetics by a continuous model. Both the SSA and the CLE capture temporal fluctuations that are due to the inherent stochasticity of the chemical process itself, often referred to as *intrinsic noise*. This intrinsic noise is attributed to probabilistic reactions occurring at low molecular concentrations. Stochastic fluctuations, however, can also stem from the interplay of the network with many other —and mostly unknown— fluctuating processes, as well as with various random physical and chemical signals arising from the extracellular world^19^. Examples arising in gene expression include the availability of promoters, RNA polymerases, the influence of long non-coding RNAs as transcriptional regulators^20^, as well as physiological differences in the cellular environment. Such sources of variability constitute *extrinsic noise*, and may cause the reaction rates of a biochemical system, and hence the firing propensity of each reaction channel, to fluctuate in time. The stochastic dynamics of a biochemical system are then attributed to fluctuations arising from both intrinsic and extrinsic noise sources — each shown to profoundly affect the system dynamics^21^. As the SSA and the CLE assume that propensities remain constant between reactions events, they cannot be used to simulate biochemical systems interacting with other networks in the cell.

A number of exact methods for simulating biochemical systems subject to both intrinsic and extrinsic noise have emerged^22–26^. While the computational efficiency of these methods have been substantially improved by use of thinning techniques, as in^23–25^, these methods still suffer similar disadvantages to the SSA: a large number of simulations may be required to obtain statistically significant results. As a result, various approximation methods have been developed, the majority of which are based upon the linear-noise approximation^27–29^. While these methods are computationally efficient, and in some cases can provide analytical expressions for the moments, they all require the assumption that the timescale of the exogenous noise process is substantially slower than the transcriptional dynamics. Some of the methods are further limited to small extrinsic noise. One exception, the Unified Coloured Noise Approximation (UCNA)^30^, allows for both slow and fast extrinsic noise timescales and leads to analytical expressions in some cases. Nevertheless, while these approaches might be an acceptable approximation in some scenarios, they completely disregard the exogenous process dynamics, and cannot be expected to hold in all timescale regimes. Indeed, external fluctuations have been shown to occur on a variety of timescales in several biological and biologically relevant settings^31–33^. For example, in^34^ it is shown that the source of the extrinsic noise (a repressor) operates on a timescale at least equal to that of the other molecules in the system. In fact, there are many instances where changes in concentration of transcription factors, that will naturally operate on a timescale comparable to the intrinsic process, result in marked biological consequences^35,36^. Any assumption of large timescale separation therefore poses potential limitations for modelling real biological systems. Furthermore, these approaches rely upon specifying a noise distribution for the parameters *a priori;* often assumed to be normal or lognormal distributions. Determining the precise extrinsic noise distributions from single-cell experimental data however is difficult, and non-identifiability issues show that the inference of such distributions is impossible beyond the most simple cases^37^.

Other approximation methods for jointly capturing intrinsic and extrinsic noise involve the incorporation of extrinsic noise into Langevin-type equations by way of *ad hoc* and often physically unfounded means. In the theory of developmental biology, for example, an extrinsic noise term is typically added to a deterministic rate equation, the form of which has been chosen to ensure the preservation of fixed points of the underlying deterministic system^4,38^. Such approaches can be justified primarily on the grounds of simplifying the mathematical analysis, but do not follow from a microscopic argument. As a consequence, the validity of these models for probing the stochastic dynamics is unclear. Since it has been shown that extrinsic noise dominates stochasticity, even in the presence of low copy numbers^31,34,39,40^, methods that formally address the inclusion of both are required.

Here we consider the chemical Langevin equation (CLE), for biochemical networks that are embedded within a dynamic fluctuating environment. The CLE, as originally formulated by Kurtz^41^, is derived directly from the underlying Markov jump process. However, we here follow Gillespie’s microscopic derivation for chemical reaction networks with time-dependent propensities, which gives rise to the *extrinsic chemical Langevin equation* (E-CLE). We show that the form of the CLE is preserved when the propensity functions are time-varying. While this may have been implicitly assumed in the literature, no formal consideration has yet been provided. Details are given in Section III. Extrinsic noise, arising as time-dependent variability in the model parameters, is introduced at the level of the CME, and the resulting E-CLE is derived from this premise: the incorporation of extrinsic noise is physically founded rather than *ad hoc*. Importantly, in Subsection III B we verify that the equations for the moments of order up to two obtained from E-CLE are exactly the same as the corresponding equations derived from the *extrinsic chemical master equation* (E-CME). In Section V, we demonstrate the accuracy and validity of the E-CLE by incorporating extrinsic fluctuations into two exemplar stochastic gene regulatory models—one linear (Subsection VA) and one non-linear (Subsection VB). For a range of different exogenous noise processes, we compare estimates of the first two moments in steady-state conditions, obtained from the E-CLE with those obtained using exact simulation methods. We further explore different timescale regimes of the extrinsic noise process and find that the E-CLE is in excellent agreement with exact stochastic simulation methods such as SSA and Extrande, as well as analytical solutions (where applicable), irrespective of the timescales considered. We show that our modelling framework offers considerable computational advantages when compared with exact methods, while still maintaining remarkable accuracy.

Finally, we show that certain limitations of the CLE carry across to the E-CLE. It is known that the CLE is unable to capture multimodality of distributions of molecule numbers in some cases. In Section VI, we extend the study of Duncan et al.^42^ to show that, while the E-CME predicts noise-induced multistability for a simple model of gene transcription, the E-CLE predicts monostability. The E-CLE is therefore not always able to capture noise induced multimodality, and should be used in conjunction with exact methods in such cases.

Altogether our results show that the extrinsic CLE framework is an efficient and accurate method for simulating the stochastic kinetics of biochemical systems subject to extrinsic fluctuations.

## II. THE CHEMICAL LANGEVIN EQUATION

Consider a well-stirred mixture consisting of *N* chemical species *S*_1_,…, *S_N_* that interact through *M* chemical reactions *R*_1_,…, *R_M_*,

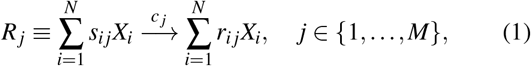

where *X_i_* denotes the number of *S_i_* molecules in the system at time *t*, and *c_j_* is the rate constant of reaction *R_j_*. Throughout, we will let **X**(*t*) = (*X*_1_(*t*),…, *X_N_*(*t*)) represent the state of the system at time *t*. Each reaction *R_j_* has an associated *propensity function a_j_* given by,

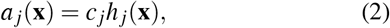

where *h_j_* (**x**) is defined to be the number of distinct combinations of *R_j_*-reactant molecules available in the state **x** (omitting here the variable *t* for brevity). The *state-change vector* **v***_j_* is defined to be the vector (*v*_1_*_j_*,…, *v_Nj_*) whose ith component is given by *v_ij_* = *r_ij_* – *s_ij_*, for *i* ∈ {1,…, *N*} and *j* ∈ {1,…, *M*}. Due to the inherent randomness of the collision of molecules in the mixture, the variable **X**(*t*) will change stochastically; mathematically, the process **X**(*t*) is a Markov jump process and can be modelled as a *homogenous Poisson process*. The time evolution of the joint probability distribution of the *X_i_* is described by the CME,

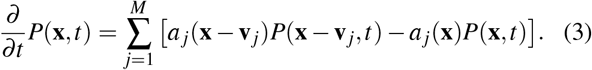

Equation (3) has a deceptively simple form: there are only a handful of systems with CMEs that are known to be solvable analytically. Simulation methods such as Gillispie’s Stochastic Simulation Algorithm (SSA) and its many variants have therefore been developed to simulate the probability distribution, *P*(**x**, *t*). While the SSA is an exact simulation method, in the sense that it generates sample paths whose probability distribution is the solution of the CME, it very quickly becomes infeasible for larger systems. In Gillespie’s paper ^17^, it is shown that under specific conditions (which can arise naturally in practise), the CME is well-approximated by a continuous Markov process satisfying the following chemical Langevin equation,

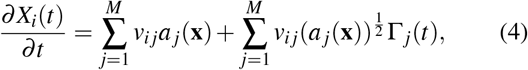

for *i* ∈ {1,…, *N*}. Here the Γ*_j_*,(*t*) are uncorrelated, statistically independent Gaussian white noises. The CLE has been shown to afford significant computational advantages compared to SSA, especially as the number and frequency of the reaction events increases.

## III. THE EXTRINSIC CHEMICAL LANGEVIN EQUATION

### A. Time-dependant propensity functions

As discussed above, biochemical systems are often subject to extrinsic sources of variability, causing the reaction rate constants, and therefore the reaction propensities of a biochemical system, to fluctuate between reaction occurrences. Both SSA and the corresponding CLE assume that the state of the system **X**(*t*), and hence the propensity of each reaction channel to fire, *a_j_*(**x**), remains constant between reaction events. Thus, SSA and the CLE capture only the random timing of reactions, and cannot be used to simulate systems whose reaction propensities fluctuate between reaction occurrences, such as those subject to extrinsic variability.

Following Gillespie^17^, we now derive the CLE for systems in which the reaction rate constants *c_j_* of Equation (1) may vary in time according to some external source. In this case, the propensity functions *a_j_* are written as *a_j_*(**x**, *t*), where the dependence on *t* is made explicit, for all *j* ∈ {1,…, *M*}, and the system **X**(*t*) is represented as *M inhomogeneous Poisson processes*. According to the random time-change representation^41,43^, the number *X_i_* of *S_i_* molecules in the system at time *t* + *τ*, for *τ* > 0, will be given by

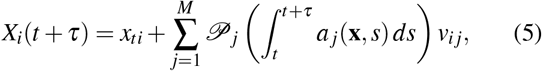

for *i* ∈ {1,…, *N*}. Each 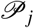 is an independent, unit rate Poisson process with rate parameter (or mean) equal to 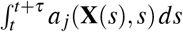, and **x***_t_* is the state of the system **X**(*t*) at time *t*. Note that the Chemical Master Equation for systems with time-varying propensities—the *extrinsic* Chemical Master Equation—is equivalent to the above Poisson formulation, and has the same form as Equation (3) above, except now *a_j_*(**x**) is replaced with *a_j_*(**x**, *t*); see the Equation A1 of the Appendix. If we further assume that the expected number of occurrences of each reaction channel *R_j_* over the duration *τ* be much larger than 1, i.e,

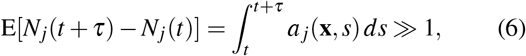

then each Poisson random variable 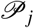 can be approximated by a normal random variable 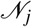 with mean and variance equal to 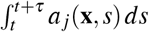. As is the case for the CLE (Equation (4)), the condition given in Equation (6) may not always be satisfied, but there are certain practical situations where this condition will hold. For small *τ*, the expected value 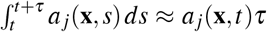, and so the satisfaction of Equation (6) will still be greatly facilitated by having large molecular population numbers. Under the condition given in Equation (6), and using the fact that a normal random variable with mean *μ* and variance *σ*^2^ can be written as 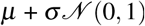, we can bring Equation (5) into the form,

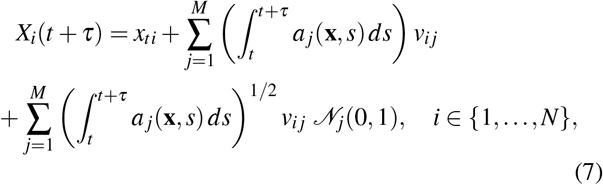

where each 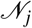 is a normal random variable with mean 0 and variance 1. Now taking the limit of (*X_i_*(*t* + *τ*) – *X_i_*(*t*))/*τ* as *τ* → 0 of both sides of Equation (7) we obtain

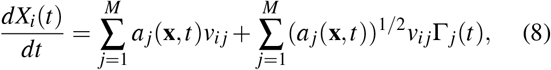

for *i* ∈ {1,…, *N*}. The Γ*_j_*(*t*) are independent Gaussian white noises (otherwise known as Wiener processes). We here used the fact that formally a Wiener process is defined as lim_*τ*→0_*N_j_*(0, *τ*^-1^) = Γ*_j_*(*t*). We refer to the CLE given in Equation (8) as the *extrinsic* chemical Langevin equation (E-CLE). For an alternative derivation of the E-CLE using the Kramers-Moyal expansion of the CME see Section A1 of the Appendix.

### B. Moments

In this section, we show that the moment equations obtained from the E-CLE and E-CME agree for moments of order up to two. As is the case for the CLE, the result is obtained from Ito’s formula^44^. We then consider what conditions are required for these equations to be solved exactly. It turns out that for systems with time-varying propensity functions, the conditions are more restrictive than for systems with constant rate parameters.

#### 1. Moment Equations

Consider an N-dimensional vector **X**(*t*) = (*X*_1_(*t*),…, *X_N_*(*t*)) satisfying the extrinsic chemical Langevin equation,

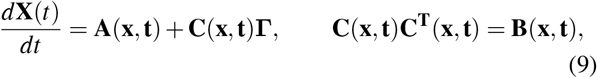

where **Γ** is a multidimensional Wiener process, and the drift vector **A** and diffusion matrix **B** are given by

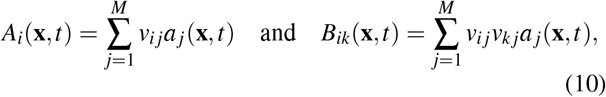

respectively. Now according to Itô’s formula, the average time-development of an arbitrary *f*(**x**) is given by

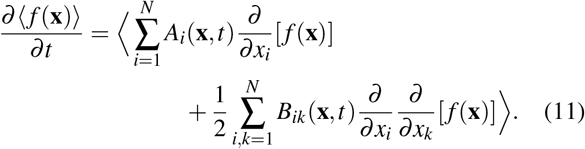

Note that here the angle bracket notation 〈·〉 denotes the expectation with respect to the solution *P*(**x**, *t*) of the extrinsic CME (E-CME) given in Eq. (A1). It then follows from Eq. (11) that for any *i* ∈ {1,…, *N*}

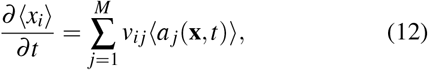

and for any *i*_1_, *i*_2_ ∈ { 1,…, *N*} we have,

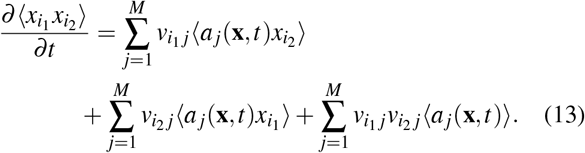

It is straightforward to show that the moment equations Eq. (12) and (13) agree with those obtained from the E-CME; the derivation involves multiplying Equation (A1) through by *x*_*i*_1__ *x*_*i*_2__ and summing over the molecule numbers. Refer to^45^ for an identical derivation of the moment equations from the standard CME. Thus, the equations for the moments of order up to two derived from the E-CME and E-CLE are identical.

It is known that the moments up to order two of the processes described by the CLE and CME agree exactly for linear reaction systems (i.e., those containing only unimolecular reactions). This is because the equations for the moments are not coupled to higher order moments, for which the evolution equations derived from the CLE and CME do not in general agree. In the case of the E-CLE, we require additional conditions to ensure that the moments up to order two agree exactly with those described by the E-CME. In particular, we require that (a) both the primary and exogenous chemical reaction networks are linear, and (b) any time-varying reaction rate parameters of the primary system occur only in zeroth-order reactions.

## IV. IMPLEMENTATION

We use the Euler-Maruyama (EM) method for simulating the E-CLE. Other implementations are possible, however, we focus on the EM method as it is the standard method for simulating the CLE. Our implementation takes as an input a pre-simulated time series of the exogenous process, typically either (i) an approximate simulation of a stochastic differential equation or (ii) an exact simulation of a chemical reaction network.

Assume we are given an input *I* that represents the exogenous process over time *t*, along with a set of coupled stochastic differential equations of the form,

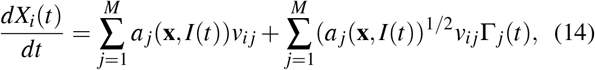

for *i* ∈ {1,…, *N*}, and where the propensity functions are time-dependent, stochastic functions that depend explicitly on the input process. The time course of the dynamic input *I* is simulated up-front; we then apply the usual EM method, but now sourcing the value of *I*(*t*) at each incremental time point of the process. This will provide an approximation **U**(*t*) to **X**(*t*) over the interval [0, *T*], conditional on the simulated noise input *I*.

1. Partition the time interval [0, *T*] into *N* equal subintervals of width *dt* = *T/N*: 0 = *t*_0_ < *t*_1_,…, < *t*_*N*–1_ < *t_N_* = *T*.
2. Initialise *U*_0_ = 0 and *I*_0_ = *I*
3. For *n* in {1,…, *N*}, recursively define

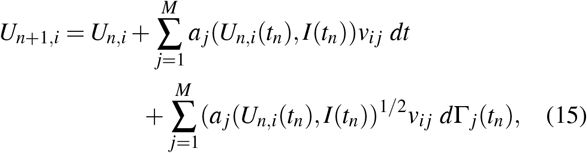

where the *d*Γ*_j_*(*t_n_*):= Γ*_j_*(*t*_*n*+1_) – Γ*_j_*(*t_n_*) are independent normal random variables with mean 0 and variance *dt*.

The function used to return the values of *I*(*t*) depends on the input process. In the case where *I* is an exact simulation of another chemical reaction network (obtained using, for example, SSA) a potential computational cost arises from the repeated search and extraction of values from the time series input *I*; this is due to the fact that SSA returns values of *I stochastically* on a discrete grid. We circumvent this by interpolating values for intermediate times by using a left endpoint constant interpolation rule. Note that no extra error is introduced by the interpolation, as it simply reconstructs the exact values of the time series at intermediate times (which are discrete, and so constant in between changes). When *I* is given by a stochastic or ordinary differential equation (SDE or ODE), *I* is simulated in parallel and synchronously with the E-CLE; thus no interpolation is required.

As a further step to improve computational performance, we note that if *I*(*t*) has not changed from the previous time increment, then we have no need at all to update the propensity functions. This occurs when the second time point in *I* is beyond the next time increment. Improvements in the runtime may be possible if, at each required sample from *I* (at time *t*_1_, say), we record the next *t*-value *t*_2_ at which the value *I*(*t*) changes. We only need to consult the list *I* again when the *t*-value has incremented sufficiently to surpass t_2_: for all intermediate values of *t*, *I*(*t*) = *I*(*t*_1_).

## V. SIMULATION

To illustrate the flexibility of the E-CLE, we consider a class of discrete transcriptional models coupled to a stochastic transcription rate. Such models have been shown to capture a range of biophysical phenomena, encapsulating empirically plausible transcriptomic distributions such as Poisson and negative binomial-like distributions, and in some cases remain amenable to mathematical analysis^46^. We focus on two important modes of gene transcription, a simple linear birth-death process, and a non-linear genetic negative feedback loop. In each case, the process is assumed to be driven by a stochastic transcription rate, *K*(*t*), which changes with time. We consider three specific dynamics^27,47^ governing the timevarying transcription rate: (1) a discrete Telegraph process, (2) a continuous Gamma Ornstein-Uhlenbeck (Γ-OU) process, and (3) a continuous Cox-Ingersoll-Ross (CIR) model. The biophysical motivations for each of these is discussed in more detail below.

We perform a comparative analysis of our method with the Extra Reaction Algorithm for Networks in Dynamic Environments (Extrande), a conditionally exact approach for the stochastic simulation of bimolecular networks with timevarying propensities^24^. Extrande is based upon thinning and rejection techniques, and has been shown to be more accurate and computationally efficient than previous methods that rely upon numerical integration of reaction propensities. We demonstrate that the E-CLE offers an accurate alternative approach that further reduces the computational time. Where feasible, we also compare our results with analytical solutions or exact simulations using Gillespie’s SSA.

Throughout, we use the following simulation parameters for the E-CLE: the time discretization of the Euler-Maruyama algorithm (*dt*), the time after which steady-state is assumed (Δ*t*), the number of samples (*N*), and the time-span for the problem (tspan). The steady-state moments are calculated from a single time trajectory by averaging over the fluctuating variables at time points Δ*t*, 2Δ*t*,…, *N*Δ*t*. To avoid break down of the E-CLE, which occurs whenever the molecule numbers become sufficiently small, we employ the complex formulation, which involves extending the domain of the E-CLE to the complex space; details of this approach can be found in^48^.

### A. Linear model: birth-death process with time-varying transcription rate

The birth-death model describes the production and degradation of a chemical species and is widely used in the study of biochemical reactions^27^. We consider a birth-death model with time-varying transcription rate *K*(*t*) as follows.

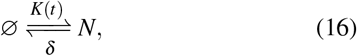

where the degradation rate *δ* is a constant parameter and *N* represents mRNA. We let the variable *X* count the number of mRNA molecules of the process described in Equation (16). Then the E-CLE (as determined by Equation (8)) for this process is given by

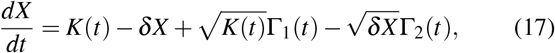

where Γ_1_(*t*) and Γ_2_(*t*) are independent Wiener processes. We refer to the birth-death part of the above *combined* process as the *primary* system, and refer to the biological processes that interact with the primary system (by way of the transcription rate) as the *secondary* or *exogenous* system.

For the remainder of this subsection, we consider two specific cases of this model more closely: (1) a discrete secondary or exogenous Telegraph system and (2) a continuous secondary or exogenous gamma Ornstein–Uhlenbeck system. These models are displayed schematically in Figure 1(a).

**FIG. 1.**
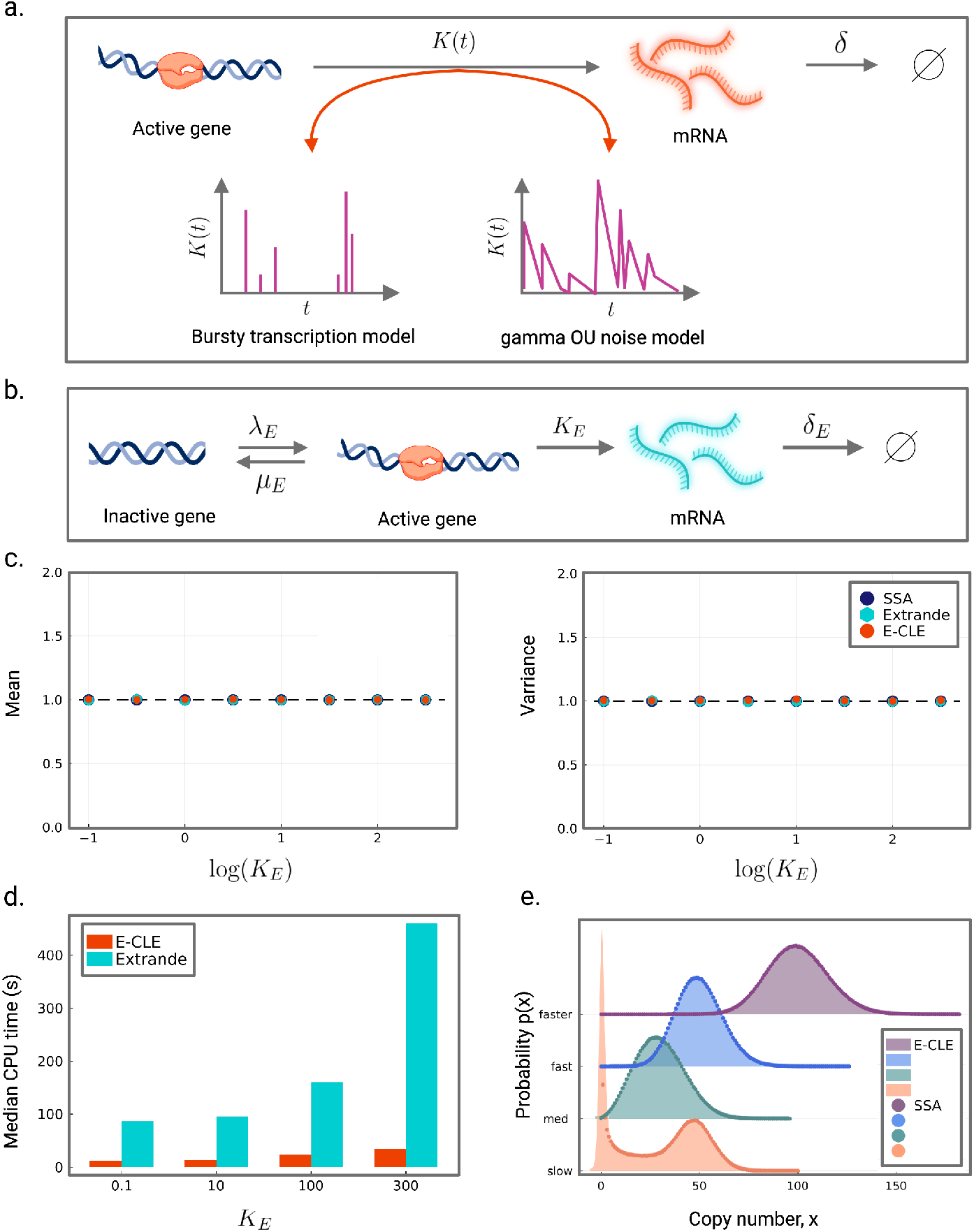
Accuracy and computational cost of the E-CLE using a linear birth-death model with time-varying transcription rate. Here *K*(*t*) is governed by a Telegraph process. (a.) A schematic of the birth-death process with time-varying transcription rate *K*(*t*). The active gene produces mRNA at rate *K*(*t*) which varies according to an exogenous Telegraph process (depicted in (b.)) or an exogenous gamma Ornstein-Uhlenbeck process (depicted in Fig. 3(a.)). The mRNA degrades at rate *δ*. (b.) The Telegraph process with active rate *λ_E_* and inactive rate *μ_E_*. The mRNA is transcribed at rate *K_E_* and degraded at rate *δ_E_*. (c.) The normalised mean (left panel) and variance (right panel) as a function of *K_E_* for the birth-death process described in (a.), where *K*(*t*) varies according to the exogenous Telegraph process in (b.) with *λ_E_* = 2, *μ_E_* = 6 and *δ_E_* = 1. The mean and variance are normalised with the exact results obtained from Gillespie simulations (shown in blue); the normalisation involves dividing the moments obtained by the E-CLE (shown in orange) and Extrande (shown in turquoise) by those obtained from the SSA. Simulation parameters used: E-CLE: tspan = 10^6^, dt = 0.01, Δ*t* = 10 and *N* = 10^5^; Extrande: tspan = 5 × 10^6^; sampling every 50 time points. The simulation parameters for Extrande are selected according to when the mean and variance converge to those of the SSA. (d.) Comparison of the median CPU time of the E-CLE (in orange) and Extrande (in turquoise) as a function of *K_E_* using the simulation parameters specified in (c.). (e.) mRNA steady-state probability distributions of the process depicted in a. obtained using the E-CLE for different promoter switching regimes of the Telegraph process. Parameters used: slow: *μ_E_* = *λ_E_* = 0.1 and *K_E_* = 50; med: *μ_E_* = *λ_E_* = 2 and *K_E_* = 60; fast: *μ_E_ = λ_E_* = 10 and *K_E_* = 100; faster: *μ_E_* = *λ_E_* = 30 and *K_E_* = 200.

#### 1. Exogenous Telegraph process

We begin by assuming the time-varying transcription rate *K*(*t*) is governed by an exogenous Telegraph process as illustrated in Figure 1(b),

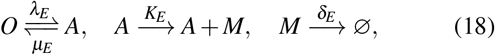

where *O* and *A* represent the inactive (“off”) and active (“on”) states respectively, and *M* represents the mRNA molecules (possibly different to the mRNA produced by the birth-death process). The parameters *λ_E_* and *μ_E_* are the active and inactive rates respectively, the parameter *K_E_* is the transcription rate, and *δ_E_* is the degradation rate. Throughout, we assume that all parameters of the combined model have been scaled by the degradation rate *δ* of the primary process, so that *δ* = 1.

Based on Section III B above, the E-CLE should predict the first two moments of the mRNA distribution of the combined process exactly. Figure 1(c) shows the mean number of mRNA molecules and the variance of fluctuations about this mean in steady-state conditions, as a function of the parameter *K_E_*. This can be viewed as varying the system size of the combined process, as the mean number of mRNA molecules of the combined process is equal to 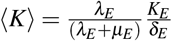, and so can be varied directly by *K_E_*. The mean and variance are normalised by the corresponding exact values obtained by stochastic simulations, using both the SSA and Extrande. The results clearly show that the E-CLE predictions for the first and second moments agree with the exact results, even for very small values of *K_E_*, corresponding to systems with very low molecule numbers.

Knowing that the E-CLE framework accurately captures mean and variance of the resulting steady-state probability density, we next compare the computational time of the E-CLE with that of Extrande. We expect the E-CLE to outperform Extrande, which relies upon thinning techniques^49^ that require two potentially computationally prohibitive steps: (1) computing propensity upper bounds of reactions and (2) an acceptance-rejection approach to handle time-dependent propensities that requires generating random numbers, see^24^ and^50^. Figure 1(d) shows the median CPU time in seconds (averaged over 5 samples) for increasing *K_E_* for both the E-CLE (turquoise) and Extrande (orange). Across all four parameter regimes, the E-CLE is considerably more efficient than Extrande. When *K_E_* is large, the E-CLE simulation is more than an order of magnitude faster. This is because an increase in the overall system size arises from an increase in the number of reactions; to achieve this increase, the time between reactions must decrease. This can be viewed as solving the system over a much finer time step, which is considerably more computationally expensive. With the E-CLE implementation, however, the time step of the Euler Maruyama algorithm is specified *a priori* and is fixed. This means that no matter how large the system is, the same number of iterations are executed over the maximum simulation time. Additionally, since the user is able to directly control the time step variable within the E-CLE framework, and with SDEs in general, it allows for even more significant speed increases (by increasing the value of the time step variable), however there is an expected trade-off with the accuracy. The E-CLE then lends itself to efficient simulation of realistic biological systems which are often large and contain multiple genes. Note that we do not include the time to simulate the secondary or exogenous process *I* since it is the same for both the E-CLE and Extrande. Additionally, our implementation of the Extrande algorithm uses the global upper bound of the pre-simulated exogenous process rather than finding local upper bounds over discretised time intervals.

To further explore the predictive capability of the E-CLE when generating steady-state probability distributions, we consider four different parameter regimes of the exogenous Telegraph process. These cover a range of promoter switching behaviours from slow to fast; refer to Figure 1(e). In the limit of slow switching between the “on” and “off” states, there is a separation of timescales between the dynamics of switching and those of birth and death of the mRNA molecules, and so the Telegraph process gives rise to a bimodal stationary probability distribution (mathematically, this is a sum of two Poisson distributions). The stationary probability distribution of the combined process is also bimodal, characterising a binary gene expression regime with high variance (Figure 1(e), orange curve). In the limit of fast switching between the “on” and “off” states, the Telegraph process gives rise to a unimodal stationary probability distribution (mathematically, this is a Poisson distribution). This is because the timescale of the mRNA is much slower than the rate of switching and will therefore approach that of a constitutively expressed gene with the same average mRNA expression levels. In this case, the stationary probability distribution of the combined process is also unimodal, with a larger-than-Poissonian variance, as depicted in Figure 1(e) (purple curve). Overall, the E-CLE closely fits the exact distributions simulated by the SSA, for all promoter switching regimes.

##### Varying environmental timescales

Considering that environmental fluctuations may operate on a range of timescales we next investigate a range of extrinsic timescale scenarios. More specifically, we consider an environmental timescale that is slow, intermediate or fast with respect to the primary (birth-death) process. The differences in speed are obtained by multiplying the vector of parameters governing the exogenous Telegraph process (namely: *λ_E_*, *μ_E_*, *δ_E_* and *K_E_*) by a constant scale parameter *s* ∈ {0.1, 1, 10} for slow-, intermediate- and fast-changing extrinsic noise regimes, respectively. Figure 2 shows the steady-state distributions of the combined process, obtained by the SSA and E-CLE. These results show that the E-CLE yields accurate distributions irrespective of the environmental timescales. We further observe that when *K*(*t*) is changing fast relative to the primary process (Figure 2 dark orange curve), the output *X* is able to suppress a significant amount of the variability in *K*(*t*). In contrast, if *K*(*t*) is changing slowly (Figure 2 blue curve), fluctuations are largely transferred to *X*. This is reflected in the variance of the distributions across the timescales, and is consistent with observations of previous studies^25^. At the limit of slow timescales in extrinsic variation, such a system approaches a mixture (or compound) model for the extrinsic noise distribution on the transcription rate for the primary model. At the other extreme, if the parameters are changing faster than the typical processes of the primary system then the copy number of the primary system will not have time to reflect the new parameter values before they have drifted again. This is much the same as a boat on choppy water – at high speeds it will experience less overall variation in height around the average than at speeds low enough to simply follow the full amplitude of the individual waves.

**FIG. 2.**
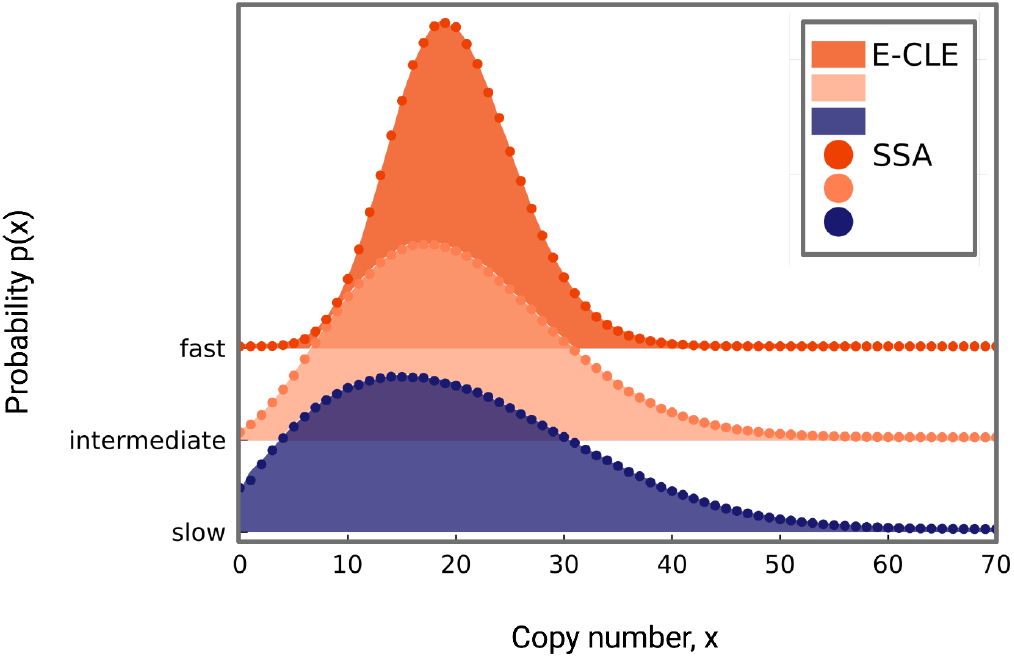
mRNA steady-state probability distributions of the combined process obtained using the E-CLE and SSA for increasing timescale separation between the primary birth-death process and the exogenous Telegraph process. The E-CLE yields accurate distributions regardless of the environmental timescale. Parameters used: *μ_E_* = 3s, *λ_E_* = 2s, *δ_E_* = *s*, *K_E_* = 50*s*, where *s* is the scaling parameter used to realise different environmental timescales. For slow *s* = 0.1; for intermediate *s* = 1 and for fast *s* = 10. E-CLE simulation parameters used: tspan = 7 × 10^6^, dt = 0.01, Δ*t* = 10 and *N* = 7 × 10^5^.

#### 2. Exogenous gamma-OU Process

We now assume that fluctuations in the time-varying transcription rate *K*(*t*) are smooth and are due to stochastic changes in the mechanical structure of DNA. Increasingly, evidence points to the fact that RNA polymerase, as it moves along the helical groove of the DNA, results in genome-wide supercoiling of the DNA during transcription^51–54^. This accumulation of mechanical stress on the DNA may obstruct the polymerases from further translocating, thereby inhibiting transcription until such time that the stress is relieved by the arrival of specific biophysical machinery known as topoisomerases. The rate of mechanical arrest is therefore fundamentally tied to the rate of mRNA production, and points to mechanical feedback as a potential source of transcriptional bursting. In order to capture the effects of these complex frustration-relaxation biomechanical properties, we follow^46^ and employ the gamma Ornstein–Uhlenbeck model, which has the following simple form,

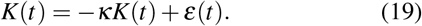

Here *κ* is the mean reversion rate representing the rate of DNA frustration, and the noise term *ε*(*t*) is a compound Poisson process of rate *a* and exponentially distributed jump sizes of rate *θ*. The parameter *a* controls the arrival frequency of topoisomerases, while *θ* controls the scale of fluctuations in the transcription rate and represents the relaxation gain. A schematic of the Γ-OU model is illustrated in Figure 3(a), where the red arrows indicate an increase in mechanical stress due to RNA polymerase-induced supercoiling and the turquoise arrows indicate stress relief by the arrival of topoisomerases.

**FIG. 3.**
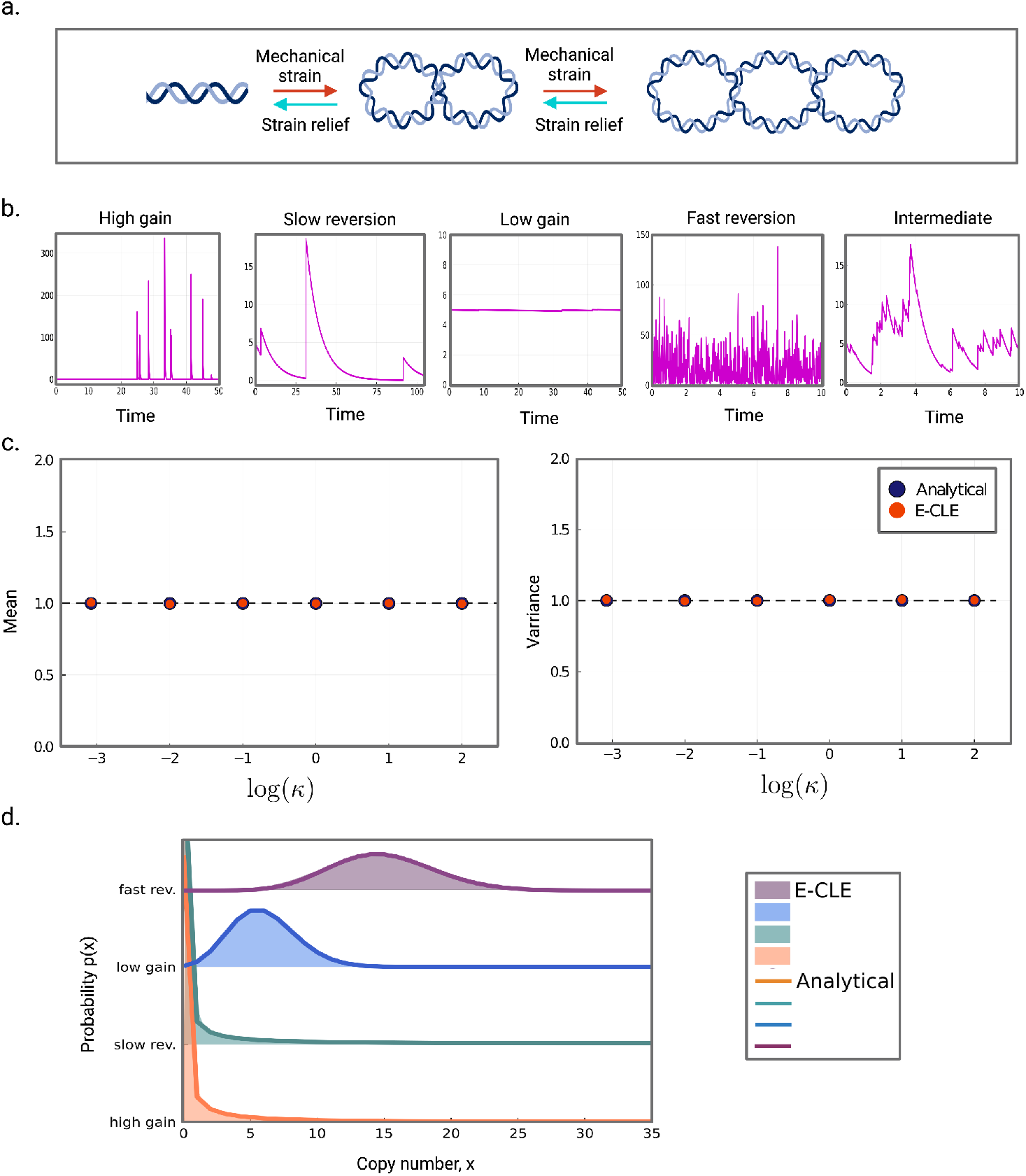
Accuracy of the E-CLE using a linear birth-death model with time-varying transcription rate. Here *K*(*t*) is governed by a gamma Ornstein-Uhlenbeck process. (a.) The gamma Ornstein-Uhlenbeck model of transcription. Transcribing DNA is placed under increased mechanical strain (shown in red arrows) with subsequent stochastic relaxation (shown in turquoise arrows) due to the arrival of topoisomerases. (b.) The transcription rate as a function of time for 5 different parameter regimes used to assess the accuracy of the E-CLE. (c.) The normalised mean (left panel) and variance (right panel) as a function of *κ* for the birth-death process in Fig. 1(a.), where *K*(*t*) varies according to an exogenous gamma Ornstein-Uhlenbeck process. The mean and variance obtained by the E-CLE (shown in orange) are normalised by the exact results, obtained analytically (shown in blue). E-CLE simulation parameters used: tspan = 10^6^, dt = 0.01, Δ*t* = 10 and *N* = 10^5^. The model parameters used are given in Table I of the Appendix. (d.) mRNA steady-state probability distributions of the combined process using the E-CLE for 4 different parameter regimes (high gain, slow reversion, low gain, fast reversion) plotted against the respective analytical solutions. Simulation parameters used: tspan 10^6^, dt = 0.01, Δ*t* = 10 and *N* = 10^5^. The model parameters used are given in Table II of the Appendix.

To explore fully the accuracy of the E-CLE when modelling steady-state probability distributions we investigate six different parameter regimes, first introduced in^46^, and outlined in Table II of the Appendix. The parameter sets cover a range of behaviours, including several limiting regimes. A representative time series depicting the typical behaviour of each regime is displayed in Figure 3(b), and can help build intuition for how each regime may affect the steady-state distribution of the combined process. In the low gain and fast reversion regimes, fluctuations either do not drift far enough from the mean value (see Figure 3(b) low gain), or revert back to the mean value so rapidly (see Figure 3(b) fast reversion) that the transcription rate may ultimately be described by the mean transcription rate of the exogenous process. Since the transcription rate is effectively constant is these cases, the combined system is expected to give rise to a Poisson distribution (see Figure 3(d) orange and green curves). In the high gain and slow reversion regimes, fluctuations can extend far from the mean (see Figure 3(b) high gain) or do not rapidly revert back to the mean value (see Figure 3(b) slow reversion), producing heavy-tailed steady-state distributions (see Figure 3(d) blue and purple curves). This intuition is formalised in^46^, where Poisson and negative binomial distributions are derived for the four limiting regimes; see^46^ for more details. In Figure 3(d), we plot the stationary probability distributions for the four limiting cases that admit analytical solutions and compare the results with those of the E-CLE. We can see that the E-CLE is an excellent match with the analytical solution in all cases. In Figure 3(d), we display the mean number of *X* molecules and the variance of fluctuations in steady-state conditions, across seven parameter regimes; these include both intermediate regimes and those displayed in Figure 3(b). The results are normalised with exact analytic results for the mean and variance obtained in^46^, which are given by,

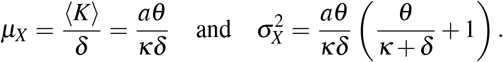

**TABLE I.** Parameter values for Figure 3(c)

| Parameter sets | $\kappa$ | $a$ | $\theta$ |
| --- | --- | --- | --- |
| Low gain 1 | $8.33 \times 10^4$ | 0.1 | 0.05 |
| Low gain 2 | 0.001 | 0.05 | 0.1 |
| Slow reversion | 0.1 | 0.01 | 15 |
| Intermediate | 1 | 4.25 | 1.493 |
| High gain | 10 | 0.1 | 150 |
| Fast reversion | 100 | 100 | 14.93 |

**TABLE II.** Parameter values for Figure 3(d)

| Parameter sets | $\kappa$ | $a$ | $\theta$ |
| --- | --- | --- | --- |
| Low gain | $8.33 \times 10^4$ | 0.1 | 0.05 |
| Slow reversion | 0.12 | 0.01 | 15 |
| Fast reversion | 100 | 100 | 14.93 |
| High gain | 10 | 0.1 | 150 |

The E-CLE predictions for the first and second moments agree with the exact results, across all regimes.

### B. Non-linear model: negative feedback loop with time-varying transcription rate

A common regulatory mechanism that arises in development is autoregulation, where transcription factors either enhance or suppress their own regulation. Here we consider a gene expression system with negative autoregulation and time-varying transcription rate *K*(*t*) as illustrated in Figure 4(a). When the gene is unbound it produces a certain protein that subsequently either degrades or binds to the active gene’s promoter. Upon binding, the active gene is rendered inactive so that no protein is produced. In this way, the protein suppresses its own production and is an example of a negative feedback loop. Such negative feedback loops have been shown to play an important role in a system’s response to noise, typically suppressing biochemical fluctuations^55^. The negative feedback system can be described by the following chemical reactions,

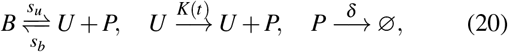

where *B* and *U* represent the bound (“off”) and unbound (“on”) states respectively, and *P* represents the protein. The parameters *s_u_* and *s_b_* are the unbinding and binding rates respectively, and *δ* is the protein degradation rate. The E-CLE (Equation (8)) for the process described in Equation (20) is then given by

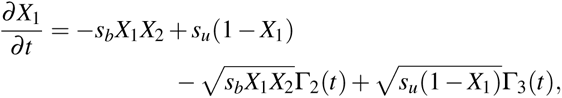

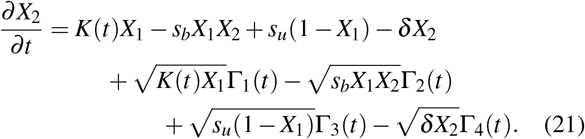

where *X*_1_ is the number of molecules of the unbound state, and *X*_2_ is the number of protein molecules. Here Γ_1_(*t*), Γ_2_(*t*), Γ_3_(*t*) and Γ_4_(*t*) are independent Wiener processes. We refer to the negative feedback loop or non-linear part of the above process as the *primary* system, and refer to the biological processes that interact with the primary system (by way of the transcription rate) as the *secondary* or *exogenous* system.

**FIG. 4.**
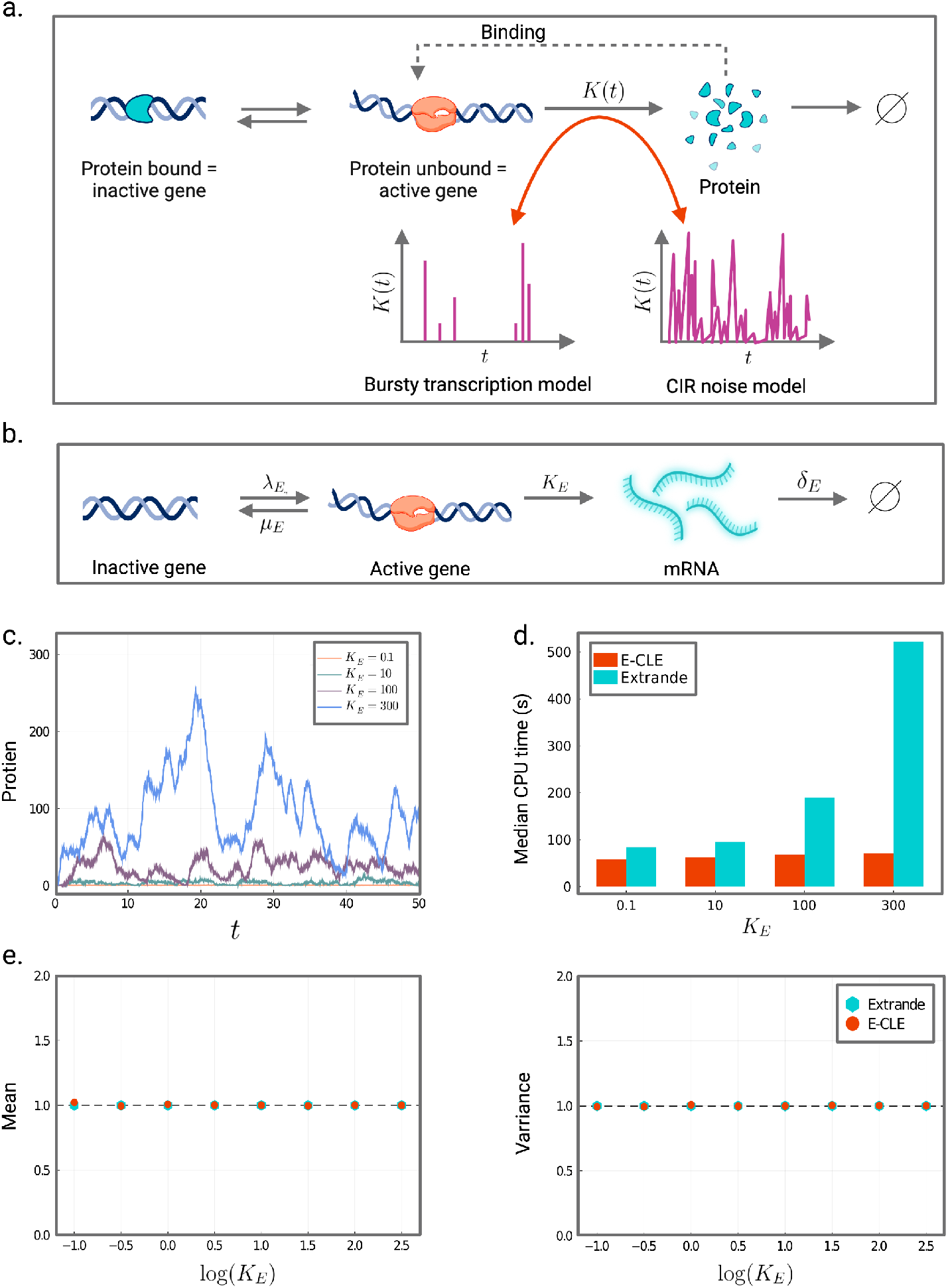
Accuracy and computational cost of the E-CLE using a non-linear genetic negative feedback model with time-varying transcription rate according to a Telegraph process. (a.) A schematic of a genetic negative feedback loop with time-varying transcription rate *K*(*t*). The active gene produces protein at rate *K*(*t*) which varies according to an exogenous Telegraph process (depicted in (b.)) or an exogenous Cox-Ingersoll-Ross process (depicted in Fig. 5(a.)). (b.) The Telegraph process with active rate *λ_E_* and inactive rate *μ_E_*. mRNA is transcribed at rate *K_E_* and degraded at rate *δ_E_*. (c.) The transcription rate *K*(*t*) as a function of time for the combined process with increasing transcription rate *K_E_* in the Telegraph process. The larger the value of *K_E_*, the bigger the system size. (d.) Comparison of the median CPU time of the E-CLE (in orange) and Extrande (in turquoise) as a function of *K_E_* on the negative feedback model subject to exogenous Telegraph noise. Simulation parameters used: E-CLE: tspan = 3 × 10^6^, dt = 0.01, Δ*t* = 10 and *N* = 3 × 10^5^; Extrande: tspan = 6 × 10^6^; sampling every 50 time points. (e.) The normalised mean (left panel) and variance (right panel) as a function of *K_E_* for the negative feedback process described in (a.), where *K*(*t*) varies according to the exogenous Telegraph process in (b.), with *λ_E_* = 2, *μ_E_* = 6 and *δ_E_* = 1. The moments are normalised by dividing the moments obtained by the E-CLE (shown in orange) by those obtained using Extrande (shown in turquoise). The simulation parameters used are given in (d.).

For the remainder of this subsection, we consider two specific cases of this model more closely: a discrete secondary Telegraph system and a continuous secondary Cox-Ingersoll-Ross process. These models are displayed schematically in Figure 4(a).

#### 1. Exogenous Telegraph process

As with the linear model, we now assume that the transcription rate *K*(*t*) of the non-linear model varies according to an exogenous Telegraph process; this is schematically illustrated in Figure 4(b.). To investigate the accuracy and computational efficiency of the E-CLE framework, we again fix the exogenous Telegraph parameters *λ_E_*, *μ_E_* and *δ_E_*, while varying the transcription rate *K_E_*, and therefore the system size. To see how the system scales with *K_E_* refer to Figure 4(c), where we plot a time series of the combined process for *K_E_* = 0.1, 10, 100 and 300. For each of the four regimes we compute the median CPU time (from five samples) using both the E-CLE and Extrande and again note that, in all cases, the E-CLE is computationally more efficient than Extrande with the highest computational gains observed for large system sizes (see Figure 4(d)). Reasoning for the reduction in computational effort follows the same logic as the linear (birth-death) model where *K*(*t*) too varies according to an exogenous Telegraph process. Lastly, we assess the accuracy of the E-CLE over eight different parameter regimes that span a range of system sizes from very few molecules to hundreds of molecules. In Figure 4(e), we plot the mean mRNA count and the variance of fluctuations about the mean, in steadystate conditions, across the different regimes using the E-CLE and Extrande. The E-CLE results are normalised with those obtained using Extrande and show that predictions for the first and second moments are in close agreement with the Extrande results.

#### 2. Exogenous Cox-Ingersoll-Ross process

Next, we assume that fluctuations in the transcription rate *K*(*t*) are smooth and depend upon the concentration of some regulatory molecule, such as RNA polymerase, or an activator or enhancer. An increase in the number of regulatory molecules, increases the frequency at which they bind to the promoter, resulting in an increase in transcription. On the other hand, a decrease in the number of molecules decreases transcription. Following^46^, we can capture this modulation in transcription using the Cox-Ingersoll-Ross model. In this model, the transcription rate *K*(*t*) evolves in time according to

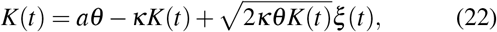

where *a* is the production rate of the regulatory molecule, *κ* is the degradation rate, *θ* is a scaling factor or regulator gain that relates the number of regulatory molecules to the rate of transcription and *ξ*(*t*) is a Gaussian white noise term.

We present a schematic of the CIR model in Figure 5(a) where regulatory molecules are shown in pink, and upon binding, enhance mRNA transcription. In Figure 5(b), we show how the system scales with the production rate *a* of the regulatory molecule. Again, we compare the computational cost of computing the stationary mRNA probability distribution of the combined process, using both the E-CLE and Extrande. Figure 5(c) shows the median CPU time in seconds (taken from five samples) for increasing values of *a*. In all cases the E-CLE (turquoise) has a lower computational burden than Extrande (orange) with more notable differences arising as the system size increases. This is because Extrande relies on computing propensities and sampling random numbers at each iteration of the algorithm, while solving the E-CLE is no more computationally cumbersome than the CLE, apart from than updating the value of *K*(*t*) at each iteration, which is solved in parallel with the E-CLE. In Figure 5(d) we show that despite a notable decrease in computation time, the mean mRNA count and the variance about this mean, computed using the E-CLE and Extrande, remain in excellent agreement for a range of parameter regimes spanning different system sizes. The E-CLE results are normalised with those obtained using Extrande.

**FIG. 5.**
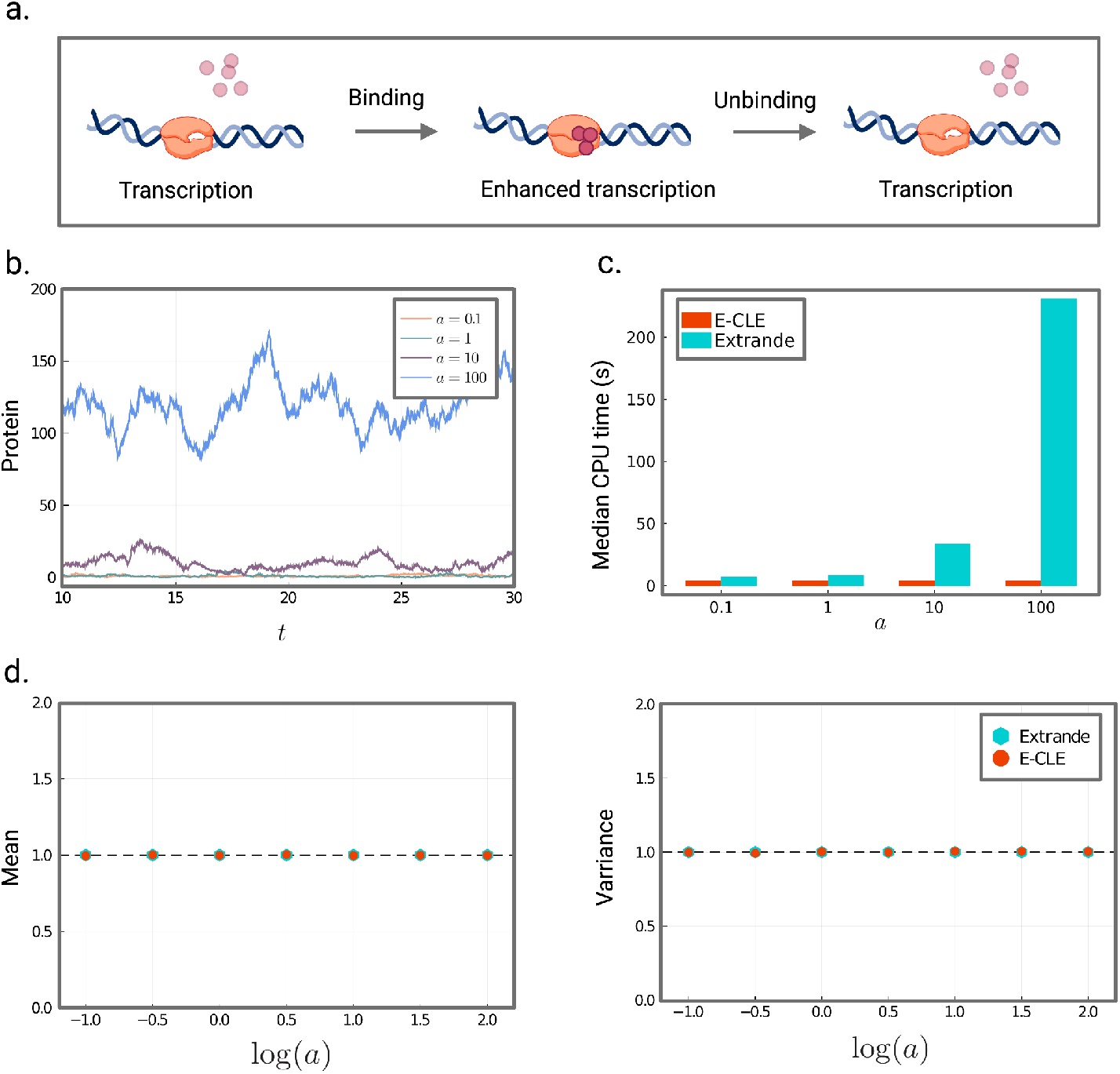
Accuracy and computational cost of the E-CLE using a genetic negative feedback model with time-varying transcription rate. Here *K*(*t*) is governed by a Cox-Ingersoll-Ross process. (a.) The Cox-Ingersoll-Ross model where transcription depends on the abundance of some regulatory molecule (show in pink). A change in the number of regulatory molecules changes their DNA binding frequency, thereby modulating transcription. Fluctuations in the transcription rate reflect fluctuations in the number of regulatory molecules. (b.) The transcription rate *K(_t_*) as a function of time for the combined process in (a.) with increasing parameter *a* in the CIR process. The larger the value of *a*, the bigger the system size. The parameters *κ* and *θ* are set to 1.25 and 10, respectively. (c.) Comparison of the median CPU time of the E-CLE (in orange) and Extrande (in turquoise) as a function of the parameter *a*. Parameters used: E-CLE: tspan = 3 × 10^5^, dt = 0.01, Δ*t* = 10 and *N* = 3 × 10^4^; Extrande: tspan = 6 × 10^5^; sampling every 50 time points. (d.) The normalised mean (left panel) and variance (right panel) as a function of *a*. The moments are normalised by dividing the moments obtained by the E-CLE (shown in orange) by those obtained using Extrande (shown in turquoise). The simulation parameters used are given in (c.).

## VI. LIMITATIONS: THE E-CLE CANNOT ALWAYS CAPTURE NOISE-INDUCED MULTISTABILITY

It has been shown in^42^ that the number of modes predicted by the chemical Fokker-Planck equation, a continuous approximation to the CME, may disagree with the number of modes predicted by the CME, in the presence of intrinsic noise. Specifically, when the CME’s marginal steadystate probability distribution changes from unimodal to multimodal, as the system size decreases and therefore intrinsic noise increases, the CFPE remains unimodal. In this way, the CFPE is unable to detect the dynamics of noise-induced multistability, defined as a change in the number of modes of the marginal steady-state probability distribution. Here we show, inevitably, that the same is true for systems modelled using the CLE subject to extrinsic noise (E-CLE). We begin by considering the model in^42^ as our primary process, but allow one of the transcription rates to vary with time according to some exogenous process:

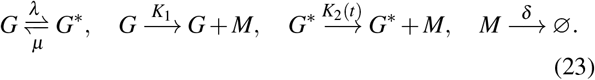

In this model, a gene is either in state *G* or *G*\*, each with a different mRNA production rate, *K*_1_(*t*) or *K*_2_(*t*), respectively. The mRNA degrades at rate *δ*, independently of the gene state. This is commonly referred to as the leaky-telegraph process since the gene is never completely inactive, but rather in a state of high production or low/leaky production. An analytical solution for the steady-state probability distribution of the leaky telegraph model was solved in^56^. As in^42^, we consider the case where there are *N* gene copies such that the total number of genes in states *G* and *G*\* equals *N* at all times (i.e., *X*_1_ + *X*_2_ = *N*, where *X*_1_ and *X*_2_ denote the number of molecules in states *G* and *G*^*^ respectively).

### 1. The E-CLE and QSA solutions disagree

Again following^42^, we consider the system described in (23) in the quasi-stationary limit (i.e., in the limit that mRNA reaches steady-state in a time much shorter than the time it takes for a gene to switch from one state to another). When *K*_2_(*t*) is constant (no extrinsic noise), the stationary mRNA probability distribution can be solved analytically as a superposition of *N* + 1 Poisson distributions; refer to^42^. We now further assume that *K*_2_(*t*) varies according to an exogenous CIR or Gamma-OU process in the low-gain or fast reversion limits. Recall that in the low gain regime, fluctuations in the underlying process barely impact the transcription rate, and in the fast reversion regime, fluctuations revert back to the mean value of *K*_2_(*t*) rapidly. Since the transcription rate is effectively constant is these cases, the limit is equivalent to a mean-field treatment of the quasi-stationary approximation (QSA) given in^42^. The stationary mRNA probability distribution can thus be written as

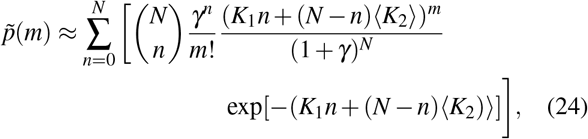

where *γ* = *μ/λ*.

We now let Ω be the compartment volume in which the chemical reaction network given in (23) is confined. Let *ϕ_m_* = *X_m_*/Ω, *ϕ*_1_ = *X*_1_/Ω and *ϕ*_2_ = *X*_2_/Ω be the concentrations of mRNA, *G* and *G*\*, respectively. Further, let *ϕ_N_* = *N*/Ω be the constant gene concentration. We will assume that *ϕ_N_* = 1, so that *N* = Ω (i.e., there is one gene per unit of volume). We now evaluate the the number of modes of the quasi-stationary distribution of *ϕ_m_* as a function of Ω. Note that the distribution of *ϕ_m_* is now given by 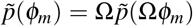. Since we have assumed the gene concentration is constant and equal to one (implying that *N* = Ω), an increase in the volume proportionally increases the total number of genes *N*, and therefore the number of mRNA molecules (this also implies there is less intrinsic noise). Thus, as the quasi-stationary distribution is a superposition of *N* + 1 Poisson distributions, we can expect 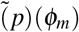 to in general be multimodal, where the number of modes increases with Ω (provided the distributions remain separated).

The E-CLE for the process described in Equation (23) can be written as

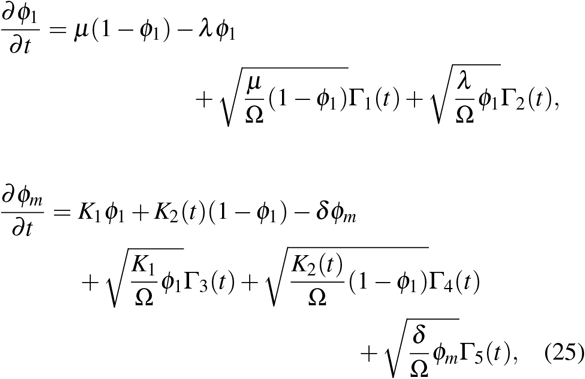

where the Γ*_i_* are independent Wiener processes, and we have used the fact that *ϕ*_1_ + *ϕ*_2_ = 1. In what follows, we will assume that all parameters of the combined model have been scaled by of the primary process, so that *δ* = 1.

In Figure 6 we plot the QSA solution in Eq. (24) and compare it with the solution from the E-CLE given in Eq (25), for increasing volumes Ω = {1, 10, 50, 100}. Following the intuition above, we see that as the volume is increased, the modality of the E-CME increases (a-c), until such a point where the modes ‘‘collide” and the quasi-stationary distribution becomes unimodal (d). The E-CLE, being a continuous approximation method, is unable to capture this multimodality since it will smooth over the two modes at *m* = 50 and *m* = 250, consequently producing a unimodal distribution with a large variance. In this way, the E-CLE is unable to predict noise-induced multimodality. We refer to the modality as noise-induced since as the volume decreases, the number of genes decreases resulting in a decrease in mRNA molecules, and as such an increase in intrinsic noise. It is important to recognise that this is an inherited limitation of the CLE, being a continuous approximation method to the CME, and not due to the addition of extrinsic noise.

**FIG. 6.**
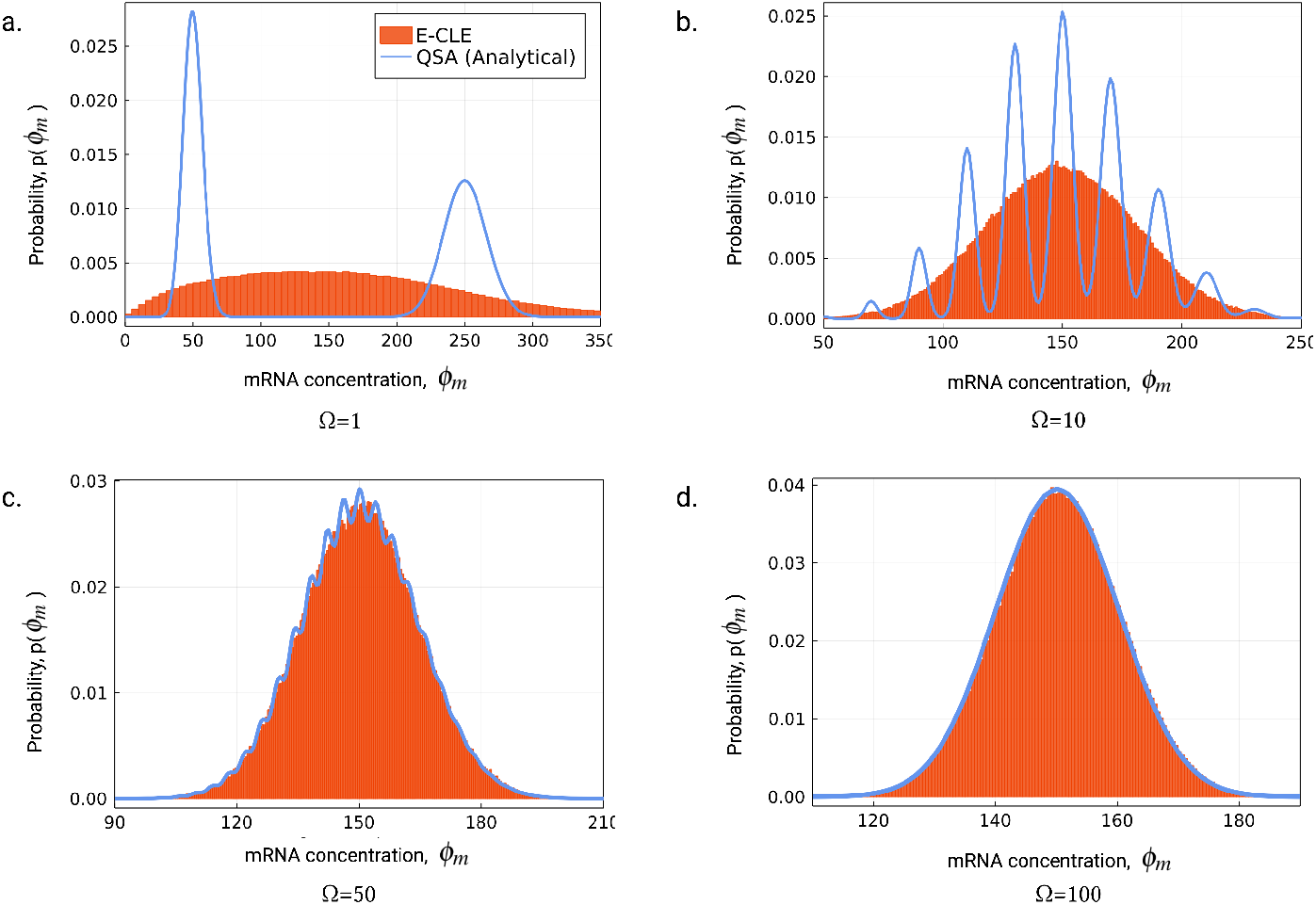
Comparison of the steady-state probability distribution of mRNA concentration *ϕ_m_* obtained from the analytical solution in Eq. (24) and the numerical solution to the E-CLE given in Eq. (25), for increasing volumes. **Ω** = {1, 10, 50, 100} in (a-d), respectively. Given that the gene concentration is constant and always equal to 1, we have *N* = **Ω**. Therefore, an increase in volume corresponds to an increase in the number of genes and subsequently an increase in mRNA molecules; and more mRNA molecules admits less intrinsic noise. Note that it is not the mRNA concentration that increases, since *N* = **Ω**, but rather the number of mRNA molecules by way of increasing *N*. Below a certain volume the QSA predicts multimodality (a-c) while the E-CLE only ever predicts one mode for all volumes (a-d). Model parameters used: *λ* = *μ* = 0.001; *K*_1_ = 50; *κ* = 8.33 × 10^-4^; *a* = 4.17; *θ* = 0.05. Simulation parameters used: tspan = 5 × 10^5^, dt = 0.01, Δ*t* = 10 and *N* = 5 × 10^4^.

## VII. CONCLUSION

Stochastic fluctuations in chemical reaction networks play a crucial role in living cells. The dynamics of such networks are modulated by both intrinsic factors, as well as those considered extrinsic, to the network such as changes in the external environment. There is now mounting biological and theoretical evidence to show that this is true for biomolecular networks in both prokaryotic and eukaryotic cells^31,34,57–59^.

Experimental approaches are becoming more sophisticated at measuring molecular variability in cell populations. The development of multi-omics for example, opens the possibility to link molecular variability between multiple regulatory layers across individual cells, and on a genome-wide scale. Theoretical and computational approaches must keep pace, if we are to model, simulate and analyse the associated data accurately. Studying the stochastic properties of gene networks is, however, challenging. It requires the correct mathematical formulation and representation of noise sources; and an efficient way to capture the resulting dynamics.

A general approach to modelling the combined effects of intrinsic and extrinsic noise is to allow the inclusion of a stochastic variable in one or more reaction rate parameters, whose dynamics may be governed, for example, by another chemical reaction network or stochastic differential equation. As a result, the corresponding propensity functions become stochastic functions of time. This renders stochastic simulation algorithms such as the SSA invalid, since the propensity functions are no longer constant between reactions events, and thus inter-reaction times are no longer exponentially-distributed. While there has been significant progress towards computationally efficient methods to simulate systems with time-varying propensities, there still remains scope for improvements. Improvements are particularly important in the context of inference problems, which are typically associated with enormous computational burden, despite algorithmic advances in the area of inference^60–63^.

Here we extend the accepted method for approximately simulating stochastic biochemical systems—the chemical Langevin equation—to include fluctuations from the surrounding environment. Our extrinsic CLE is derived directly from the CME meaning that fluctuations, both internal and external, are physically founded. We view this as safer than including *ad hoc* or heuristic terms, which are prone to corrupting the downstream analysis. Importantly, we show that the equations for the moments up to order two of the processes described by the E-CLE and E-CME are identical.

The E-CLE offers an accurate and efficient framework for probing the stochastic dynamics of biochemical systems subject to both intrinsic and extrinsic noise. The E-CLE is found to be accurate even for chemical systems with species in very low molecule numbers, for both the linear and non-linear system studied above. We demonstrate that the predicted distributions and moments of the E-CLE are robust under a range of noise processes and time-scale scenarios. Further, we show that even using a standard Euler-Maruyama implementation and employing the complex formulation^48^ (which typically requires more samples to achieve accurate estimates than other implementations), the E-CLE is computationally advantageous compared to the conditionally exact simulation algorithm Extrande. From a modelling perspective, the E-CLE is appealing because the equations are easy to derive and implement for simulation.

Despite the generality of the E-CLE, it is not without caveats. First, the E-CLE suffers similar disadvantages to the CLE: (i) it breaks down in finite time whenever molecule numbers become sufficiently small and (ii) its solution may not always agree with that of the discrete E-CME, particularly in the case of noise-induced multi-stability. The former restriction can mostly be rectified, however, by employing the complex formulation, the CCLE, which has been shown to restore the accuracy of the CLE in such cases^48^. A second limitation, and in practical terms arguably minor, is that the E-CLE assumes that the state of the primary system does not influence the extrinsic noise process, allowing the input into the E-CLE to be pre-simulated. The E-CLE should be applicable to a variety of extrinsic noise sources, including those arising from differences in cell cycle duration^19,57^, cell size^64–66^ and cell shape^67^, as well as those arising from the physical environment such as temperature, pressure or light.

We expect the accuracy and speed gains introduced by the E-CLE to be even more pronounced for larger chemical reaction systems, e.g. whole cells^68^. Increased computational speed can enable us to study new types of scientific problems. The speed increases found in Fig. 1, for example, have the potential to introduce a step-change in the system size that we can study.

## DATA AVAILABILITY STATEMENT

All models and simulations were implemented in The Julia Programming Language (v.1.6.3). Code implementing an example of the E-CLE is publicly available at https://github.com/theosysbio/extrinsic_CLE.

## ACKNOWLEDGMENTS

The authors gratefully acknowledge David Schnoerr for helpful discussions throughout this research. L.H. and M.P.H.S. are supported by the University of Melbourne DVCR Driving Research Momentum fund. M.A.C. is supported by a University of Melbourne graduate research scholarship.

## Appendix A: Appendices

### 1. Kramers-Moyal Expansion

Here we present an alternative derivation of the E-CLE using the Kramers-Moyal expansion of the CME truncated to the first two terms. The derivation closely follows that of the standard CLE. Our starting point is the following Chemical Master Equation, which is equivalent to the random time-change representation given in Eq. (5).

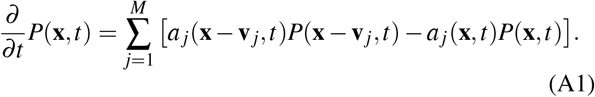

Taking the first term of the right-hand side of Eq. (A1), and performing a Taylor expansion to the second order around ***x*** we get,

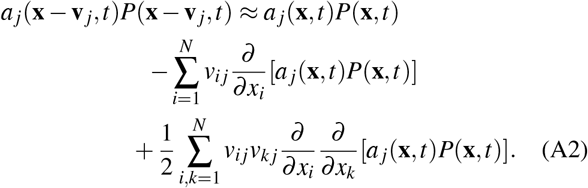

Substituting this equation into the CME, we see that the first term on the right-hand side of Eq. (A2) cancels with the last term on the right-hand side of the CME. Thus, the CME simplifies to

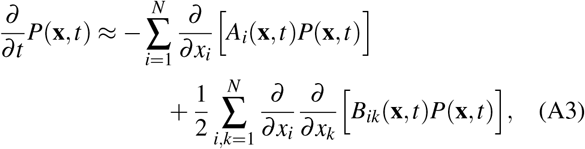

where **A**(**x**, *t*) is the drift vector and **B**(**x**, *t*) is the diffusion matrix, defined respectively by

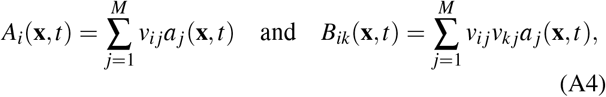

Note that both **A**(**x**, *t*) and **B**(**x**, *t*) depend on time due to the fact that the propensity functions depend on time. Eq. (A3) takes the form of a Fokker-Planck Equation (FPE) which is equivalent to the Itô stochastic differential equation^47^

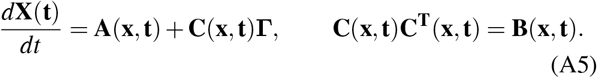

Here **Γ** is a multidimensional Wiener process. Now choosing *C_ij_* = *v_ij_*(*a_j_*(**x**, *t*))^1/2^, we arrive at the E-CLE (given in Eq. (8)),

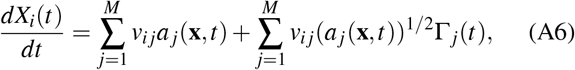

for *i* ∈ {1,…, *N*}.

### 2. Parameter regimes for gamma-OU noise

The following parameter regimes are used in the simulation of the birth-death process with gamma-OU noise (Subsection VA).

